# Potential of Negative Ion Mode Proteomics: MS1-Only Approach

**DOI:** 10.1101/2023.03.06.530802

**Authors:** Pelayo A. Penanes, Vladimir Gorshkov, Mark V. Ivanov, Mikhail V. Gorshkov, Frank Kjeldsen

**Affiliations:** Department of Biochemistry and Molecular Biology, University of Southern Denmark, DK-5230 Odense M, Denmark; V. L. Talrose Institute for Energy Problems of Chemical Physics, N. N. Semenov Federal Research Center for Chemical Physics, Russian Academy of Sciences, 38 Leninsky Pr., Bld. 2, Moscow 119334, Russia

## Abstract

Current proteomics approaches rely almost exclusively on using positive ionization mode, which results in inefficient ionization of many acidic peptides. With an equal quantity of acidic and basic proteins and, correspondingly, the similar number for their derived peptides in case of the human proteome, this inefficient ionization poses both a substantial challenge and a potential. In this work, we study the efficiency of protein identification in the bottom-up proteomic analysis performed in negative ionization mode, using the recently introduced MS1-only ultra-fast data acquisition method DirectMS1. This method is based on accurate peptide mass measurements and predicted retention times. Our method achieves the highest rate of protein identifications in negative ion mode to date, with over 1,000 proteins identified in a human cell line at a 1% false discovery rate using a single-shot 10-min separation gradient, which is comparable with hours-long MS/MS-based analyses. Evaluating the proteins as a function of pI indicated preferable identification of the acidic part of the proteome. Optimization of separation and mass spectrometric experimental conditions facilitated the performance of the method with the best results in terms of spray stability and signal abundance obtained using mobile buffers at 2.5 mM imidazole and 3% isopropanol. The work also highlighted the complementarity of data acquired in positive and negative modes: Combining the results for all replicates for both polarities, the number of identified proteins increased up to 1,774. Finally, we performed analysis of the method’s efficiency when different proteases are used for protein digestion. Among the four studied proteases (LysC, GluC, AspN, and trypsin), we found that trypsin and LysC performed best in terms of protein identification yield. Thus, digestion procedures used for positive mode proteomics can be efficiently utilized for analysis in negative ion mode.

---

Mass spectrometry (MS)-based proteomics facilitates characterization and quantitation of the proteins present in an organism, including modifications, structures, and functions. This analysis can reveal detailed information about sub-cellular protein location, protein-protein interactions, and protein structural dynamics. Additionally, quantitative proteomics can help researchers better understand disease mechanisms, cellular functions, and discover potential biomarkers.^1,2^ In a standard large-scale proteomics study, electrospray ionization (ESI)–MS in positive ion mode in conjunction with reversed-phase HPLC is the preferred technique for identifying tryptic peptides.^3^ This includes using acidic modifiers (typically formic acid) in the mobile phases, providing solid protonation conditions. However, this approach might not be best suited for analyzing many acidic peptides, which tend to lose protons upon ionization. This can potentially render the analysis suboptimal for approximately half of the human proteome, in which acidic and basic proteins and their derived peptides are roughly equally present.^4,5^ Furthermore, many post-translational modifications (such as phosphorylation^6^, glycosylation,^7^ and sialylation^8^) carry acidic functional groups, thus, suggesting preferable ionization as anions.

Hence, ESI–MS in the negative mode should be more suited for optimal ionization of acidic peptides, and has been reported to produce less background noise compared with the positive one.^9^ Nevertheless, using negative polarity in proteomics is not as explored as the positive one due to several technical difficulties. The first limitation of negative ion mode is the presence of corona discharge,^10^ which affects spray stability and degrades instrument performance. Several approaches have been suggested to solve this issue: (i) external gas flow (SF_6_ or dry O_2_) perpendicular to the column^11^, (ii) fluorinated and chlorinated solvents to work as electron scavengers^12^, (iii) organic modifiers (e.g., methanol or isopropanol) in the mobile phases to reduce the dielectric constant of the eluent^13,14^, and (iv) mechanically assisted ionization methods [capillary vibrating sharp-edge spray ionization (cVSSI)].^15,16^

In addition, collision-induced dissociation (CID) in the negative ion mode is complicated by internal fragmentation and many neutral losses (e.g., H_2_O and NH3) that render spectra difficult to interpret and predict.^17^ Although Cassady^18,19^ and Bowie^20,21,22^ have made efforts to understand the fragmentation patterns of individual peptides using CID, application on a proteome-wide scale has not been demonstrated yet. Electron detachment dissociation^23^ (EDD) is an alternative to collision– activation fragmentation by using fast, high-energy (> 10 eV) electrons to detach electrons from deprotonated anions, leading to spontaneous peptide fragmentation. Although EDD is of limited efficiency, its fragments are mainly dominated by *a*- and *x-* ions.^24^ Other more-adaptable fragmentation techniques for large-scale proteomics studies include negative electron detachment dissociation (NETD)^25,26,27^ and negative ultraviolet photodissociation (NUVPD).^28,29^ These two fragmentation techniques have provided substantial gains to the figures of merit in negative ion mode proteomics, and can be adapted with dedicated instrumental and operational modifications.

All the aforementioned fragmentation techniques are MS/MS methods that include scheduled events of precursor ion isolation, accumulation, activation, and product ion detection. Despite continuous MS improvements, such event adds substantial time to the total instrumental cycle time. A simpler and potentially efficient approach to negative in mode proteomics would be a methodology that refrains from the MS/MS event and relies on other peptide characteristics for confident protein identifications. In such a case the proteomics experiment would no longer depend on product ion yield, predictability of the fragmentation pattern, or even time spend in the MS/MS. Instead, simple detection of even low-abundance peptide ions in the MS 1 scan eluting from the chromatographic system would be sufficient for assessing the proteome. Even though most common proteomics studies depend on MS/MS data, reliable protein identification can be obtained without fragmentation.^30,31^ Several methods for MS 1-only proteome analysis have been developed for positive ion mode, ranging from the early accurate mass tag^32^ (AMT) method to more recent approaches that rely on the accuracy of peptide *m/z* measurements and retention time (RT) predictions. RTs are especially important for MS1-only strategies, as they contain the only sequence-specific information in the (*m/z*, RT)- space. With the development of machine learning algorithms, highly accurate RT prediction models are becoming increasingly available^33,34^. The DirectMS1 method, which we previously described,^30^ provides false-discovery (FDR) controlled protein identification and quantitation based on proteolytic peptide analysis using high-resolution MS acquired in an MS1-only mode of instrument operation. Briefly, the method is built around the *ms1searchpy* search engine^30^ employing advanced machine learning algorithms and tools for identifying proteins present in the sample based on detected peptide-like LC-MS features. Without the MS/MS event, meaningful depth of quantitative proteome coverage was demonstrated within a few minutes of experimental time. Specifically, DirectMS1 identified up to 2,000 proteins from mammalian cell lines using a 5-min HPLC gradient (in positive ion mode).^35^ Such throughput can be particularly relevant in research that involves a large number of samples, such as clinical population studies.^36^ The number of MS1 spectra that can be processed in this manner is only determined by the acquisition rate of the mass analyzer operating at high mass-resolution settings. Our aim in this study is to capitalize on previous experience and develop the first MS1-only methodology for conducting pro-teome-wide analysis in the negative ion mode.

## EXPERIMENTAL SECTION

### Samples

All chemicals were purchased from Sigma–Aldrich (St. Louis, MO, USA) unless otherwise stated. Optimization of the chromatographic method was performed using *E. coli* cell lysate and the final assessment of the method was performed using *HeLa* cell lysate. Measurements for the highest methodology performance were performed using Thermo Scientific Pierce *HeLa* protein digest standard (P/N 88328) derived from the *HeLa* S3 cell line.

### Sample preparation

*E. coli* cells were prepared as follows: an aliquot of 10^6^ cells was resuspended in 200 μL lysis buffer [6M guanidinium chloride, 10 mM tris(2-carboxyethyl)phosphine (TCEP) (Thermo Scientific, Rockford, IL, USA), 20 mM chloroacetamide (CAA), and 100 mM triethylamonium bicarbonate (TEAB) buffer; pH 8]. Cells were incubated in a shaker at 300 rpm at 90 °C for 30 min. Cells were lysed using an ultrasonic homogenizer (QSonica sonicators equipped with a microprobe, Newton, CT, USA) for 2 min with 90% amplitude on ice, with 1-s on/off pulses. A 50-μL aliquot was then diluted with 250 μL of 100 mM TEAB pH 8 prior to digestion with trypsin at a ratio of 1:50 w/w and incubated at 37 °C for 4 h. After this time, a trypsin boost at a ratio of 1:100 w/w was added, and the digestion continued at 37 °C overnight. The reaction was quenched by adding formic acid (FA; Merck, Darmstad, Germany) to a final volume of 1% v/v. For the *HeLa* cells, an aliquot of ≈10^6^cells was resuspended in 400 μL lysis buffer (1% w/w SDC, 10 mM TCEP, 20 mM CAA, and 100 mM TEAB buffer; pH 8) and cells were incubated in a shaker at 300 rpm at 90 °C for 30 min. Cells were lysed using an ultrasonic homogenizer (QSonica) for 4 min with 40% amplitude on ice, with 1-s on/off pulses. Prior to digestion, an 80-μL aliquot was diluted with 120 μL of TEAB buffer; 100 mM, pH 8. Each of the enzymes [trypsin (Promega, Madison, WI, USA), GluC (Roche, Mannheim, Germany), LysC (Promega, Madison, WI, USA) and AspN (Merck, Burlington, MA, USA)] were added at a ratio of 1:50 w/w and incubated for 4 h;at 25 °C in the case of GluC and 37 °C for the rest of the enzymes.At this time, a booster of each enzyme at a ratio of 1:100 w/w was added to the corresponding digest, and the digestion was performed at 25 °C for GluC and 37 °C for the rest of the enzymes overnight. The reaction was quenched by adding FA to a final volume of 1% v/v. After sample digestion and before HPLC–ESI–MS analysis, both *E. coli* and *HeLa* samples were desalted using Sep-Pak Vac 1cc (50 mg) C18 cartridges (Waters, Milford, MA, USA). Once the samples were desalted, they were subsequently dried and redissolved with 100 μL of Milli-Q water, and the concentration of each sample was measured using the Qubit protein assay (Thermo Fisher Scientific).

### LC optimization

LC methods were optimized by using both LTQ Orbitrap XL and Q Exactive HF mass spectrometers (Thermo Fisher Scientific, San Jose, CA, USA) coupled with an UltiMate 3000 LC system (Thermo Fisher Scientific, Germering, Germany). A μ-Precolumn C18 PepMap100 trap column (5 μm, 300μm i.d., 5mm length, 100 Å; Thermo Fisher Scientific, USA) and a self-packed analytical column (XBridge Premier Peptide BEH C18, 2.5μm, 75μm i.d., 18cm length, 130 Å; Waters, Milford, MA, USA) were used for the separation. In each injection, 1 μg of the *E. coli* digest and a mixture of iRT peptides (Biognosis AG, Schlieren, Switzerland) at 1× concentration were injected. Mobile phase A consisted of the corresponding modifier (FA in positive mode, and imidazole/piperidine in negative mode) in Milli-Q water and mobile phase B consisted of 95% ACN (VWR Chemicals, Rosny-sous-Boix, France) in water with the modifier at the same concentration as in mobile phase A. For the sake of simplicity and readability, the concentration of the modifier will be specified in each of the experiments. When isopropanol was present in the mobile phase (1% to 5%), it was added at the same concentration in both mobile phases. The loading solvent was in all cases mobile phase A. The LC gradient was from 2% to 40% B in 60 min at a flow rate of 250 nL/min.

### Training of the DeepLC model for negative ion mode

For training the retention time prediction, a 15-μg aliquot of the trypsin and GluC *HeLa* digest was loaded onto an Acquity UPLC M-Class CS C18 column (Waters, Etten-Leur, Netherlands) for offline reversed-phase low pH fractionation using a Dionex Ultimate 3000 HPLC system (Thermo Scientific, Bremen, Germany). Peptides were loaded using buffer A (0.1% FA in water) and separated by increasing concentrations of buffer B (95% ACN and 0.1% FA in water) from 2% to 40% B in 70 min and from 40% to 95% B in 5 min. After 10 min at 95% B, mobile phase composition was switched back to 2% B in 1 min and was held at 2% B for 4 min. Peptides were collected in 30 concatenated fractions in 10 different wells, using 3 min per eluting fraction. Fractions were dried and redissolved in buffer A before LC analysis. Fractions were subsequently analyzed using an Orbitrap Fusion Lumos mass spectrometer (Thermo Scientific, San Jose, CA, USA) coupled to an UltiMate 3000 LC system (Thermo Fisher Scientific, Germering, Germany). All data for the training was acquired in DDA mode in positive mode using alkaline mobile phases. MS/MS identifications were mapped to the LC-MS features detected by *biosaur2* utility^37^ and then used to train the model.

### Data acquisition for method evaluation

Final LC–MS/MS analysis for method evaluation was performed using an Orbitrap Eclipse mass spectrometer (Thermo Scientific, San Jose, CA, USA) coupled to an UltiMate 3000 LC system (Thermo Fisher Scientific, Germering, Germany). For these short gradients (5, 10, and 15 min), a self-packed trap column (XBridge Premier Peptide BEH C18, 2.5μm, 150-μm i.d., 3cm length, 130 Å; Waters, Milford, MA, USA) and a self-packed analytical column (XBridge Premier Peptide BEH C18, 2.5μm, 75-μm i.d., 5-cm length, 130 Å; Waters, Milford, MA, USA) were used. The mobile phases were as follows: i) positive ion mode: (A) 0.1% FA in H_2_O and (B) 0.1% FA and 95% ACN in water; ii) negative ion mode: (A) 2.5 mM imidazole and 3% IPA in water and (B) 2.5 mM imidazole, 3% IPA, and 95% ACN in water. The loading solvent was in both cases mobile phase A. The gradient was from 2% to 40% B in the corresponding minutes of the gradient (5, 10, or 15 min) at a flow rate of 1.5 μL/min. Full MS scans were acquired with 240,000 resolution. For the Thermo Scientific Pierce *HeLa* protein digest standard, 500 ng of sample was injected. In the case of the digestion with the different enzymes, 1 μg of *HeLa* digest was loaded on the column. For each condition (positive and negative), each chromatogram time (5, 10, and 15 min) and each enzyme (trypsin, GluC, LysC, and AspN), the sample was injected in quadruplicate.

### Data analysis

Raw files were converted into mzML format using ThermoRawFileParserGUI (version 1.7.0). Peptide features matches (PFMs) were detected using *biosaur2*^37^(version 0.2.3) and the following parameters: minimum number of points to define a peak=1, maximum *m/z1500*. Proteins were identified by using *ms1searchpy* (version 2.3.6), which is the search engine of DirectMS1^35^ (freely available at https://github.com/markmipt/ms1searchpy under Apache 2.0 license). The parameters used for the search were the following: a minimum of one scan for the detected isotopic cluster, a minimum of one visible C13 isotope, charges from 1–6 (both in positive and negative mode), no missed cleavages, and an initial mass tolerance of 8 ppm. All searches were performed against the Swiss–Prot human database containing 20,185 protein sequences concatenated with decoys^38^. Results were filtered to 1% FDR at the protein level. Joint results were obtained by combining standalone ms1searchpy searches using *ms1combine_proteins* script distributed with *ms1searchpy*. Briefly, standalone protein search scores were summed, and proteins were re-filtered to 1% FDR using the target-decoy approach in the same manner as in the standard *ms1searchpy* search.

Protein pI distributions were calculated using the *Peptides* R package^39^ using the Lehninger scale. Protein filtered lists were analyzed with DAVID^40, 41^ for enrichment of cellular compartment analysis using the *Homo sapiens* proteome as background.

## RESULTS AND DISCUSSION

When using negative mode polarity in ESI, both rapid needle deterioration and reduced efficiency of ionization can be caused by the phenomenon of corona discharge. This occurs at the tip of the ESI emitters, where free electrons are accelerated in the high-field region produced during ESI. This can lead to a decrease in spray stability and a loss of ion signal. To improve the ESI stability, we adjusted first the quantity of isopropanol (IPA) added to the mobile phase. We examined six concentrations of IPA (from 0% to 5%) in both mobile phases (A: 0.1% FA in water; B: 95% ACN + 0.1% FA in water). Our results are in line with previous observations^14^ and indicate that higher concentrations of IPA led to a more stable spray, and the voltage required to achieve this stability was also lower as the IPA concentration increased. Additionally, increasing IPA concentrations also reduced the retention times of the iRT peptides (sequences provided in Table 1 of Supplementary Information), with hydrophilic peptides being more affected than hydrophobic peptides (Figure 1a). The influence of IPA on the ion abundance was also evaluated, and for this purpose we measured the same iRT peptides in the matrix of an *E. Coli* tryptic digest, in both positive and negative mode. The results (Figure 1b) indicate that IPA had no measurable negative effect on the ion signal abundance.

**Figure 1.**
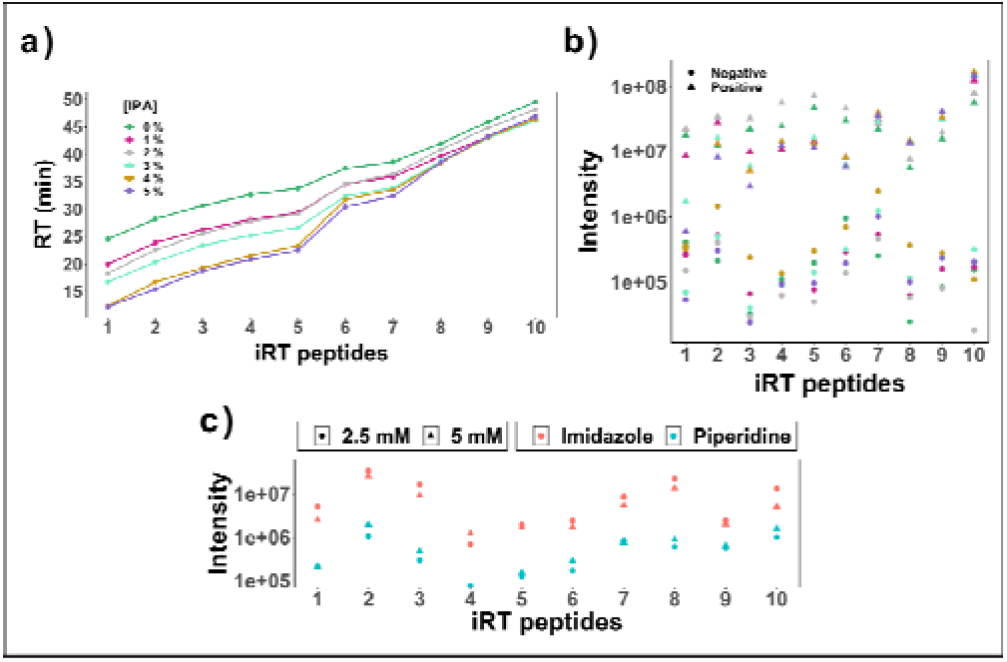
Chromatographic optimization: (a) Variation of peptide retention time as a function of IPA concentration; (b) Peptide ion abundance as a function of IPA concentration for both polarities; (c) Peptide ion abundances using imidazole and piperidine (2.5 and 5 mM).

Advancement in negative ion mode proteomics also requires conditions that result in the highest possible anion flux. Without appropriate solvent modifiers the difference between positive and negative ion mode is roughly two orders of magnitude. Various solvent systems have been proposed and used with great success, in particular various concentrations of piperidine.^28, 42, 43^ We also speculated that imidazole could function as an attractive modifier in negative ion mode, since imidazole has an aromatic ring where the positive charge can be efficiently delocalized between both nitrogen atoms present in the molecule. Similar stabilization is not possible with piperidine, as the ring only has one nitrogen atom and no aromaticity, which renders imidazole an intrinsically stronger base. Hence, we compared both piperidine and imidazole in concentrations of 2.5 and 5 mM. The results in Figure 1c indicate that the peptide ion abundances were generally greater with imidazole than piperidine as the modifier. A concentration of 2.5 mM was the more suitable than 5 mM in the mobile phase because the pH in the mobile phase does not change substantially (≈9 and 9.2 for 2.5 and 5 mM, respectively), whereas the salt concentration in the mobile phase is double, which can compromise spray formation.

After this critical evaluation, we selected 2.5 mM imidazole and 3% IPA in both mobile phases as the most suitable conditions for negative ion mode analysis.

We used a *HeLa* standard as a reference sample to evaluate the performance of the MS1-only negative ion mode regime. Figure 2a shows the average number of identified proteins in both modes, indicating comparable results between positive and negative mode; with more than 700, 1,000 and 1,200 identified proteins for the 5, 10, and 15-min gradients, respectively. To more thoroughly evaluate the data, we focused on and inspected the 10-min gradient data, as it presents a good compromise between protein identification yield and chromatogram length. By combining the data of all four replicates in each mode, a Venn distribution (Figure 2b) shows the number of identified proteins in positive (1556) and negative (1259) modes. This leaves 10% of additional proteins present only in negative mode (162). If identified proteins from each polarity are represented as a function of the isoelectric point (pI) (Figure 2c) and compared with the theoretical distribution, it is possible to observe a preference for the acidic part of the proteome recorded in the negative mode and *vice versa*.

**Figure 2.**
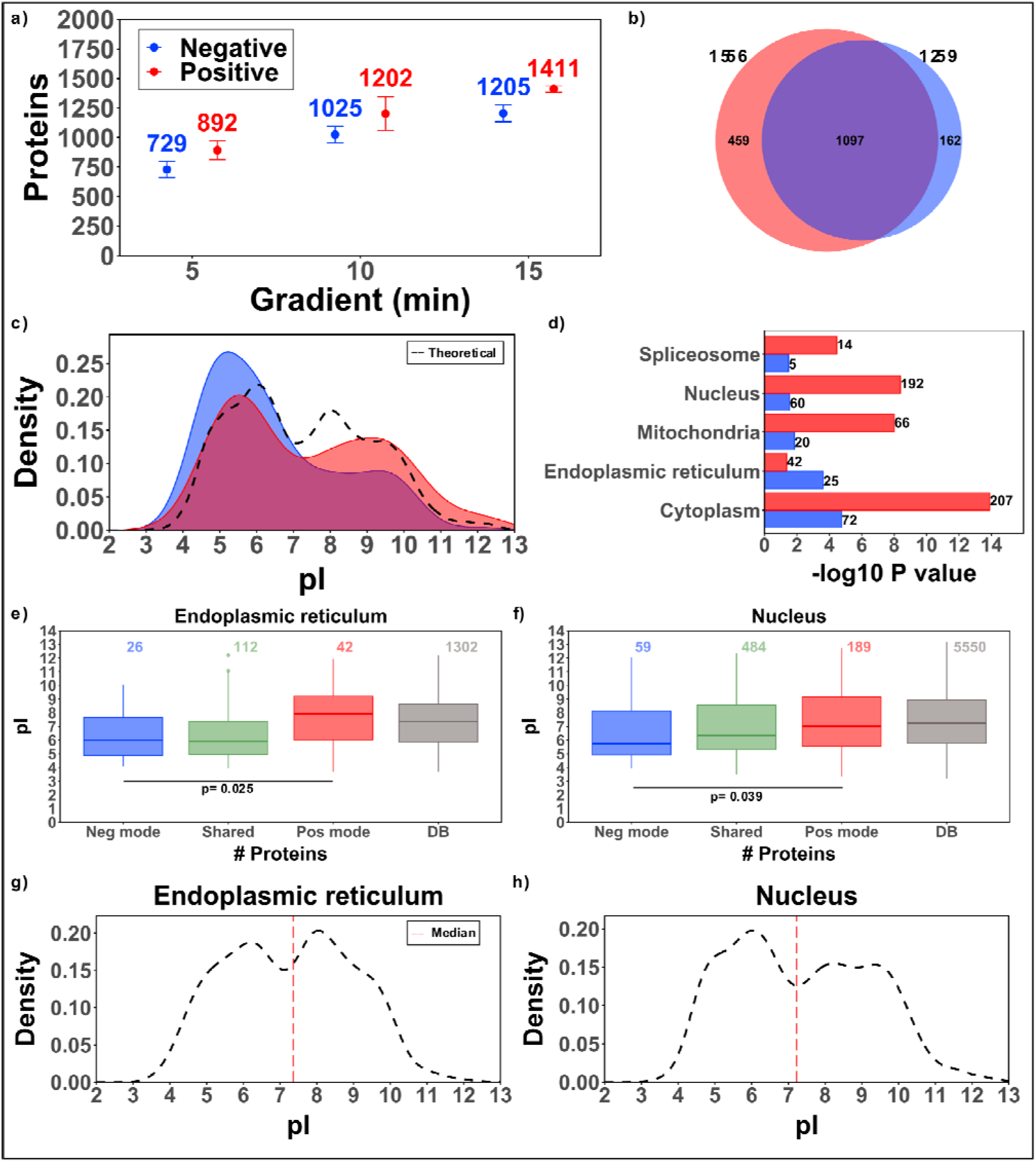
Evaluation on *HeLa* standard data: (a) Average number of identified proteins for the different gradients in each mode (error bars indicate the standard deviation of four replicates); (b) Venn diagram of the identified proteins for the combined replicates with the 10minute gradient; (c) Protein distribution as a function of pI in each mode, and compared to the theoretical distribution; (d) -log P value of the most enriched subcellular compartments of the identified proteins unique to either positive (red) or negative (blue) mode; (e) Boxplot of the pI distribution of the identified proteins present in the endoplasmic reticulum displayed in accordance with the polarity and compared with the protein database distribution; (f) Boxplot of the pI distribution of identified proteins present in the nucleus displayed in accordance with the polarity and compared with the protein database distribution; (g) Database protein distribution of endoplasmic reticulum proteins as a function of pI (median value highlighted in red); (h) Database protein distribution of nucleus proteins as a function of pI (median value highlighted in red).

The pI of a protein is a principal characteristic that defines the net charge of a protein at a specific pH. Properties of the proteome-wide pI distributions have been studied and found to be bimodal, with the majority of proteins exhibiting a pI value of *ca*. 5.0 and 9.0.^44^ The relationship between protein pI and subcellular localization has also been analyzed and indicates that there are strong statistically significant correlations between protein pI and subcellular localization. For example, mitochondrial and nuclear locations have a higher propensity to comprise more basic than acidic proteins; whereas the opposite is true for lysosomal, cytoskeletal, peroxisomal, and cytoplasmic locations. This suggests that the variation of localization-specific pI distributions might be linked to local pH differences in subcellular compartments.^45^ Based on this knowledge we investigated if proteins from different cellular compartments would tend to become enriched by using different ionization modes. Figure 2d shows the topmost enriched cellular compartments (spliceosome, nucleus, mitochondria, endoplasmic reticulum, and cytoplasm) when proteins found only in either positive or negative ion mode were considered. Both modes share cytoplasmic proteins as first rank, but second rank are nucleus and endoplasmic reticulum proteins for positive and negative mode, respectively. Considering this, we divided the proteins for the combined 10-min gradients in three different subsets as in Figure 2b: positive (459), negative (162), and shared (1097) (the list of proteins is provided in Table 2 of the Supplementary Information); and we compared each subset with the protein distribution from the reviewed SwissProt database for the nucleus and endoplasmic reticulum. Boxplots in Figure 2e and 2f clearly indicate that negative ion mode has a higher preference for identifying acidic proteins compared with positive mode for both endoplasmic reticulum and nucleus proteins. In fact, this trend is consistent for all five cellular compartments investigated (Figure S1). Contrary to expectation, the protein pI distribution of the nucleus or endoplasmic reticulum (Figure 2g and 2h) is neither particularly acidic nor basic, and cannot account entirely for the order of the enriched cellular compartments.

Previous studies have demonstrated the value of using multiple proteases in both positive and negative ion mode bottom-up proteomics.^46, 47^ Although the potential of using different enzymes has been documented for MS1-only approaches in positive ion mode,^37^ this possibility has not been studied for negative ion mode. Hence, we digested *HeLa* cells using trypsin, LysC, GluC, and AspN; with the aim of investigating the suitability of various proteases for DirectMS1 experiments, as well as to enhance the coverage and depth of the proteomic analysis. The results in Figure 3a show the average of the four replicates for every enzyme and gradient at the protein level. Most interestingly, LysC and trypsin yield the best results, both in positive and negative ion mode. Our initial assumption was that GluC and AspN would generate the most suitable peptides for analysis in the negative mode, since the cleavage site of these enzymes ensure at least one acidic residue in each peptide. Although unexpected, the remarkable performance of trypsin and LysC suggests that standard digestion protocols used in typical positive ion mode proteomics can be applied without modification in negative ion mode analyses as well. Although AspN and GluC performed best in negative mode compared with positive mode, protein identification for these enzymes could not match the results from LysC and trypsin.

**Figure 3.**
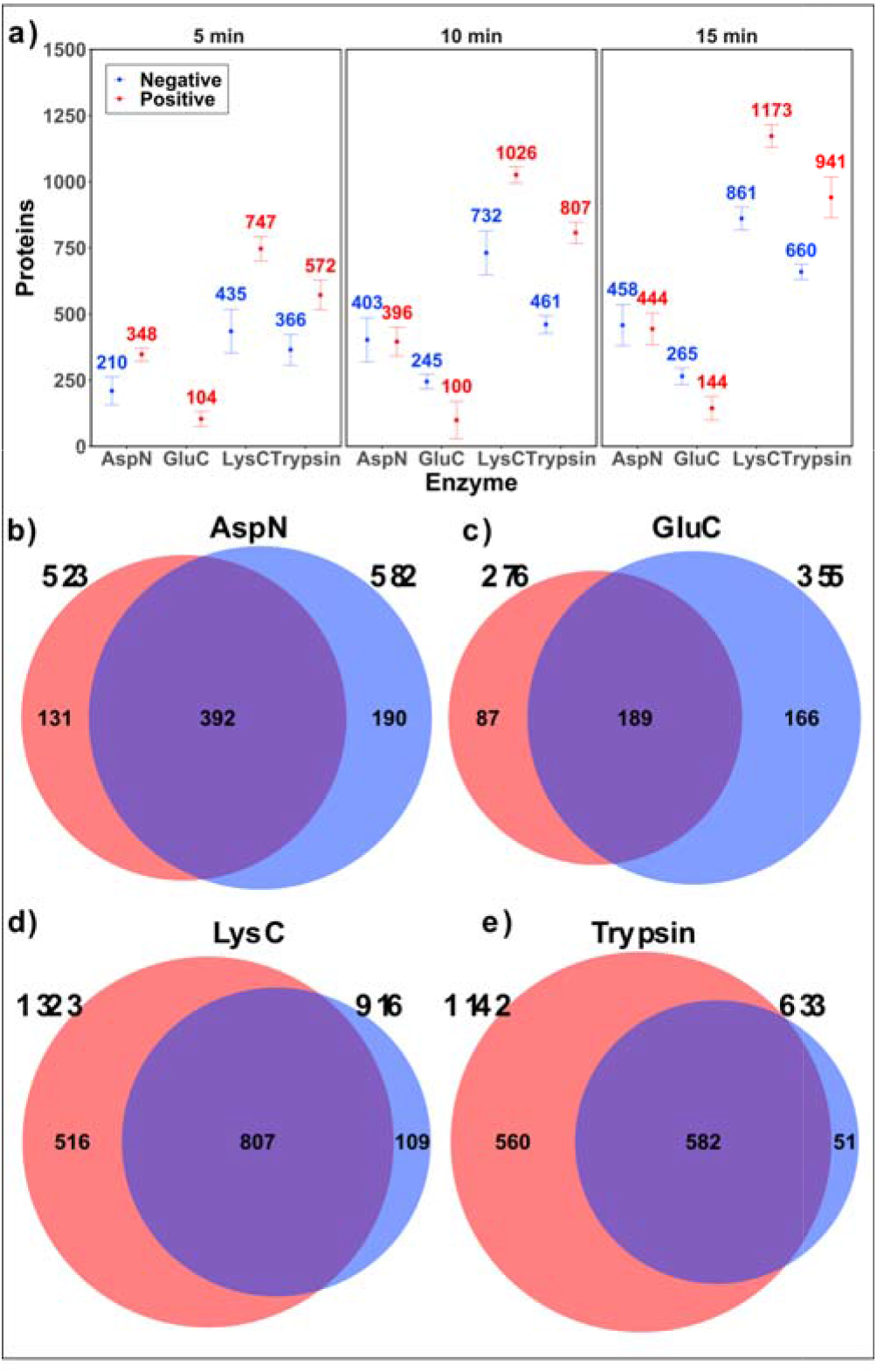
Evaluation of *HeLa* digest: (a) Average number of identified proteins for the different gradients and enzymes in positive and negative mode (error bars indicate the standard deviation of the four replicates); Venn diagrams of the identified proteins (negative in blue; positive in red) using replicates from 10-min gradient and either (b) AspN; (c) GluC; (d)LysC; or (e) trypsin as proteases.

As in the case for the *HeLa* standard, we combined the four replicates from each mode (10-min gradients), and Venn diagrams in Figure 3b–3e show the number of identified proteins for each enzyme. These results indicate the complementarity of using different enzymes in negative ion mode; with *ca*. 10% new proteins in negative mode for AspN, LysC, and trypsin. In the case of AspN and GluC, more than 33% and 60% of the identified proteins in negative mode were not identified in the positive mode, respectively. Despite this large difference, the number of identified proteins remains the lowest when compared with the other the enzymes.

To understand why trypsin and LysC were the best matches to negative ion mode, we focused on the charge state of the detected PFMs. Every PFM might be in more than one charge state, and for each PFM, we counted the state with the highest abundance. These results are shown in Figure 4a and 4b for negative and positive ion mode, respectively. Whereas doubly charged PFMs were the most abundant for all four enzymes in both modes, singly charged PFMs were almost nonexistent in the positive mode. In the negative ion mode, peptides of trypsin and LysC digestion had the highest prevalence of singly charged PFMs, accounting for 22.8% and 15.6% respectively; whereas AspN and GluC had a lower occurrence of singly charged PFMs, only accounting for 7.7% and 9.9%, respectively. The DirectMS1 methodology has a distinct advantage in that it can gain PFM information even from singly charged peptides. In contrast, most MS/MS fragmentation techniques (and particularly electron-based fragmentation techniques such as ECD/ETD, EDD, and NETD) rely on multiple charged peptides for successful peptide sequencing. Besides being efficient proteases, the substantial contribution to the PFM pool from singly charge peptides renders trypsin and LysC attractive as proteases for this approach in negative ion mode proteomics.

**Figure 4.**
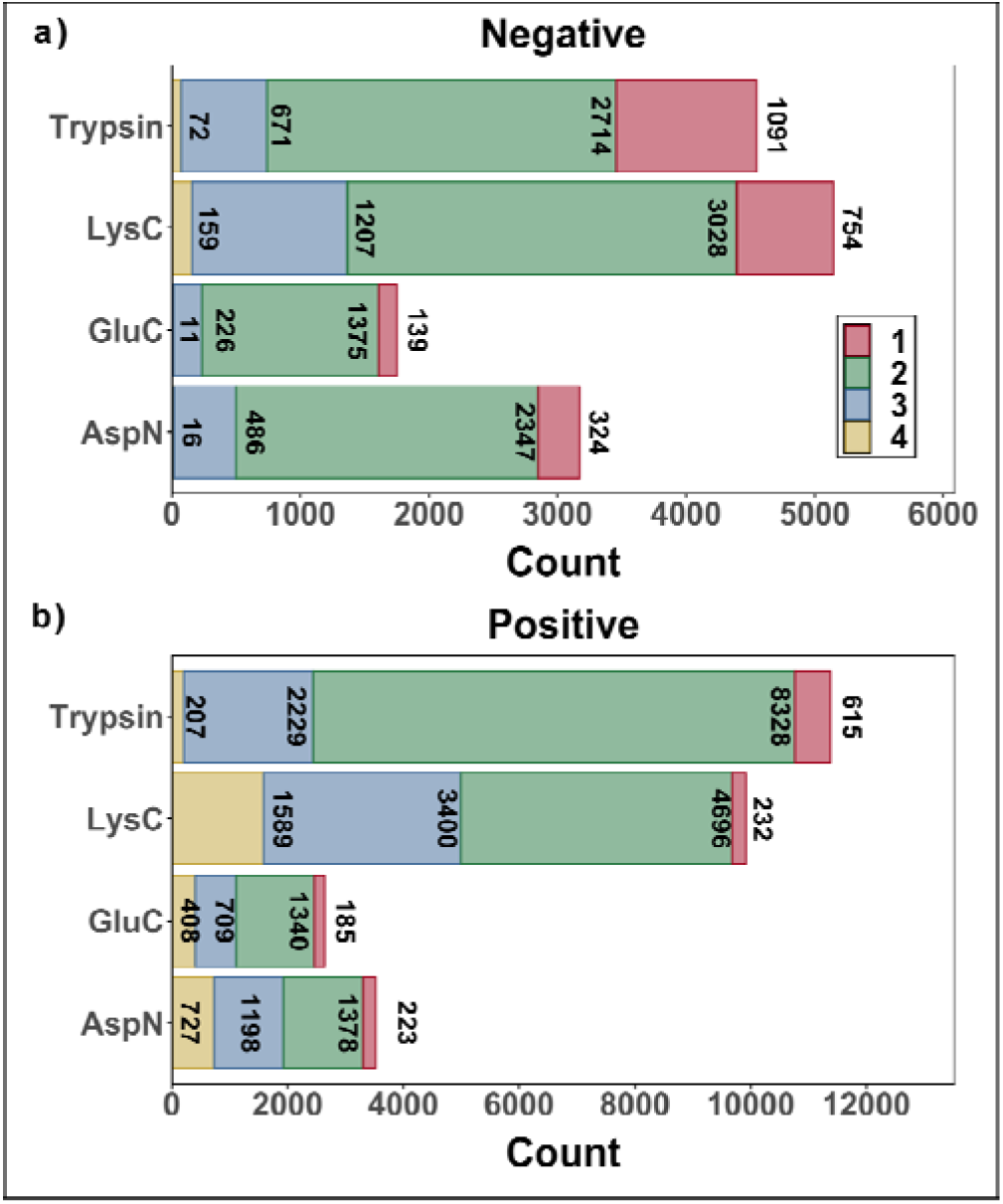
Charge state distribution of the *HeLa* digest: Most abundant PFM charge state for (a) negative ion mode and (b) positive ion mode data for each protein digest.

### Combined data search

So far, we have analyzed the performance of protein identification in both ion modes separately and discovered several benefits. Merging the PFM features from both ion modes prior to protein scoring could offer even greater advantages, especially when only a limited number of PFMs are found in both modes. Individually, the data from either ion mode might not be sufficient to reach a significant protein identification score. However, combining the PFM data from both modes can lead to a higher score if there is substantial complementary information between the two modes. Specifically, we assume such integration to yield a general higher sequence coverage per protein as well as an increase in the number of confidently identified proteins. The results in Figure 5a for the *HeLa* standard indicate that for all gradients, the combination of the data from both polarities resulted in a higher number of identified proteins, with 1774 proteins for the 10-min gradient and more than 2000 proteins for the 15-min gradient data. In the case of the different enzymes (Figure 5b–5e), a combined data search generated higher number of identified proteins except for the data for trypsin and 10-min gradient data. This can be explained by the poor statistics or casuistry of the results. Evaluation at 5% FDR rate (data shown in Table 2 of the Supplementary Information) confirms that protein identification is higher when using data from both polarities compared to the data acquired only in positive/negative mode. These results further underscore the complementary of two ionization modes in proteomics studies.

**Figure 5.**
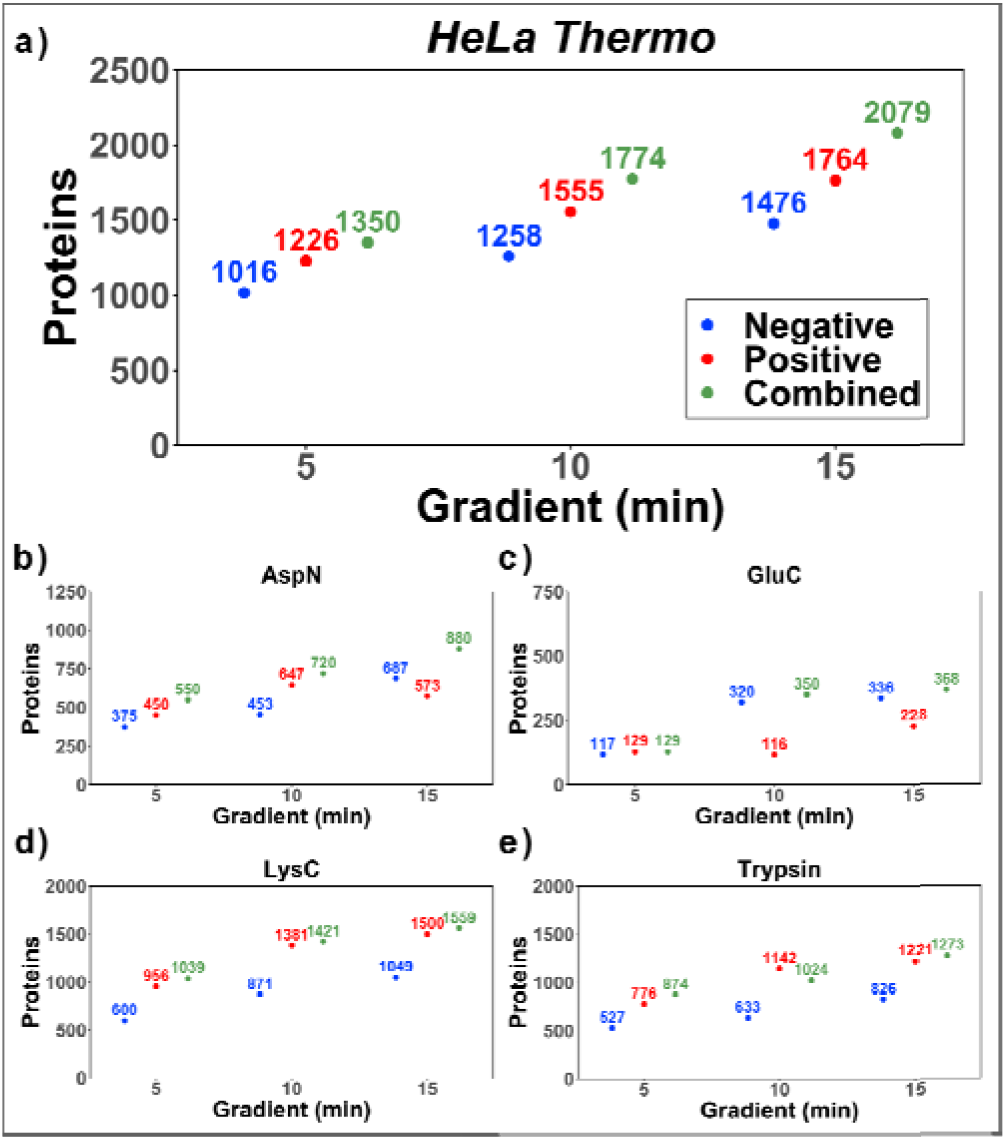
Number of identified proteins for the combined search of all replicates in positive, negative, and both modes, for: (a) *HeLa* standard (trypsin); (b) *HeLa* AspN digestion; (c) *HeLa* GluC digestion; (d) *HeLa* LysC digestion; (e) *HeLa* trypsin digestion.

## CONCLUSIONS

Our study has provided a comprehensive evaluation of negative ion mode ESI-MS for proteomic analysis, using anMS1-onlyapproach. Through optimizing the chromatographic conditions, we achieved reproducible chromatography and high ionization efficiency, resulting in similar protein identification compared with that of the positive ion mode. Moreover, a joint search of the data acquired in both modes afforded an increase in the number of identified proteins to 1774 and 2079 proteins for the 10- and 15-min gradient data, respectively. Accordingly, negative ion mode proteomics provides complementary sequence and proteome coverage, with a gain biased toward the acidic part of the proteome. Additionally, our results suggest that standard digestion protocols used in typical positive ion mode proteomics (trypsin and LysC digestion) can be applied without modification for negative ion mode proteomics. DirectMS1 has the advantage of providing highly efficient protein quantification^48^ and being independent of the charge state of peptide precursor ions. We underlined the latter point by reporting trypsin and LysC as the best-performing proteases even in the negative ion mode, despite substantial formation of many singly charged peptides. Further efforts will focus on evaluating the quantitative capabilities of using joint data from positive and negative ion mode.

## Supporting information

HeLa standard data. Unique protein ID negative mode

HeLa standard data. Unique protein ID positive mode

HeLa standard data. Protein ID shared between pos. neg mode

## Author Contributions

F.K conceptualized and supervised the study;F.K, P.A.P, and V.G. refined and planned the experiments, P.A.P. and V.G. performed experiments; and M.V.I. developed the search algorithm. P.A.P, V.G., M.V.I, M.V.G., and F.K analyzed the data; P.A.P and F.K. wrote the manuscript with input from all the authors. All authors discussed the results and commented on the manuscript. All authors have given approval to the final version of the manuscript.

## Notes

The authors declare no competing financial interest.

## DATA AVAILABILITY

The data sets generated and analyzed during the current study have been deposited to the ProteomeXchange Consortium via the PRIDE partner repository with the dataset identifier PXD040583.

## ACKNOWLEDGMENT

We would like to thank Dr. Robbin Bouwmeester for his valuable input for the negative ion mode DeepLC model retraining. This study was supported by the Lundbeck Foundation (Grant R346-2020-1215 to F.K.). Proteomics and mass spectrometry infrastructure at the University of Southern Denmark SDU was supported by generous grants to the VILLUM Center for Bioanalytical Sciences (VILLUM Foundation grant no. 7292), PRO-MS: Danish National Mass Spectrometry Platform for Functional Proteomics (grant no. 5072-00007B), and the Novo Nordisk Foundation (INTEGRA, NNF20OC0061575). *ms1searchpy* development was supported by the Russian Science Foundation, grant no. 21-74-10128.

