## Supplementary information for "Potential of Negative Ion Mode Proteomics: MS1-Only Approach"

**Supporting materials**

**Description of the joint search**. Joint results were obtained by combining standalone ms1searchpy searches using ms1combine_proteins script distributed with ms1searchpy. Briefly, standalone protein search scores were summed, and proteins were re-filtered to 1% false discovery rate using target-decoy approach in the same way it is done in the standard ms1searchpy search. However, in such a simple summing approach, all proteoforms in protein groups may be reported, while standalone searches prevent such situations due to the rule used in the search “one peptide scored only for one protein”. When we have multiple search results, there are small chances that peptides will be used for scoring different proteoforms in different runs. In that case, the sum of scores for all proteoforms may be above the FDR threshold and all of them will pass it. To prevent such situations, we choose the best file from all experimental data in terms of the number of identified proteins at 1% FDR. The list of all proteins from the database for this chosen file was taken and these proteins were ranked using their ms1searchpy scores. After that, protein scores for all ms1searchpy runs were recalculated using the principle “one peptide scored only for one protein with best rank”. Thus, any peptide shared between proteoforms was scored only for a single proteoform for all experimental data used in this project.

Basically, the described results could be obtained using three steps:

1. Standard ms1searchpy search.

Command: ms1searchpy somefile1.mzML …(some options).

2. Protein score recalculation.

Command: ms1combine somefile1_PFMs_ML.tsv -fdr 1.0 -pp path_to_bestfile_proteins_full.tsv -out somefile1_RECALC.

3. Final joint results.

Command: ms1combine_proteins somefile1_RECALC_proteins_full.tsv … somefileN_RECALC_proteins_full.tsv -fdr 1.0.

**Supplementary Table 1**: Sequences of the synthetic iRT peptides used for chromatography and mass spectrometric optimization.

| Peptide number | Sequence |
| --- | --- |
| 1 | GAGSSEPVTGLDAK |
| 2 | VEATFGVDESNAK |
| 3 | YILAGVENSK |
| 4 | TPVISGGPYEYR |
| 5 | TPVISGAPYEYR |
| 6 | DGLDAASYYAPVR |
| 7 | ADVTPADFSEWSK |
| 8 | GTFIIDPGGVIR |
| 9 | GTFIIDPAAVIR |
| 10 | LFLQFGAQGSPFLK |

**Supplementary Table 2**: Excel files containing the list of proteins detected in the HeLa standard sample.

| Mode | File name |
| --- | --- |
| Positive | Only_positive.csv |
| Negative | Only_negative.csv |
| Both modes | Shared.csv |

Supplementary Table 3: Protein identification for the combined data at 5% FDR for all positive and negative replicates, as well as the combined search (positive and negative mode simultaneously).

|  | **Polarity** | | |
| --- | --- | --- | --- |
| **Enzyme** | **Positive** | **Negative** | **Both combined** |
| *HeLa* Thermo | 1982 | 1611 | 2242 |
| AspN | 846 | 745 | 1041 |
| GluC | 407 | 511 | 685 |
| LysC | 1781 | 1225 | 1838 |
| Trypsin | 1527 | 930 | 1571 |

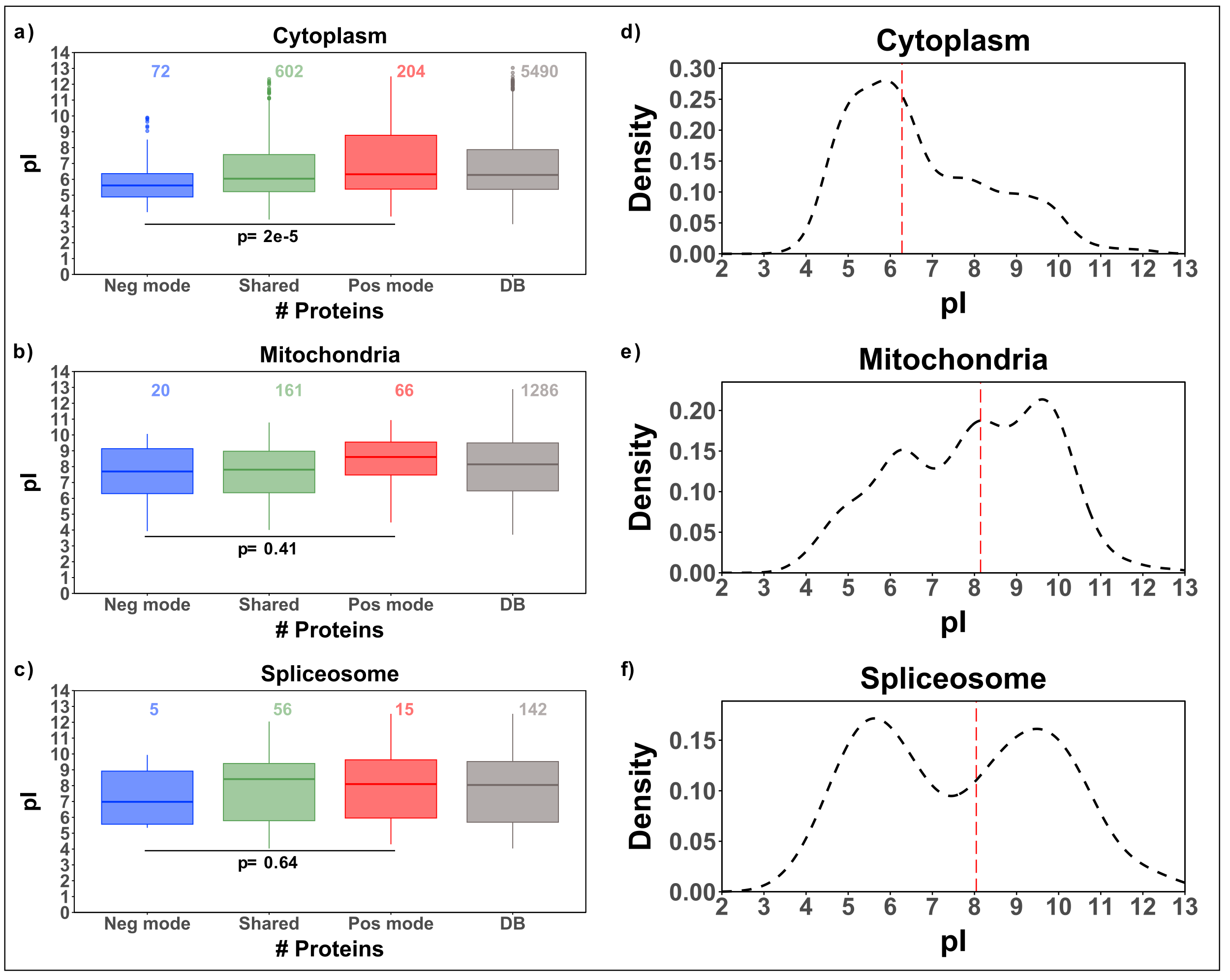

**Supplementary Figure S1**: (a) Boxplot of isoelectric point (pI) of the identified proteins present in the cytoplasm separated according to the polarity and compared to the proteome distribution; (b) Boxplot of pI of the identified proteins present in the mitochondria separated according to the polarity and compared to the proteome distribution; (c) Boxplot of pI the identified proteins present in the spliceosome separated according to the polarity and compared to the theoretical distribution; (d) Proteome distribution of cytoplasm proteins as a function of pI; (e) Proteome distribution of mitochondria proteins as a function of pI; (f) Proteome distribution of spliceosome proteins as a function of pI.
